# Structural basis for no spectral shift of heliorhodopsin by counterion mutation

**DOI:** 10.1101/2020.06.24.168591

**Authors:** Tatsuki Tanaka, Manish Singh, Wataru Shihoya, Keitaro Yamashita, Hideki Kandori, Osamu Nureki

## Abstract

Microbial rhodopsins comprise an opsin protein with seven transmembrane helices and a retinal as the chromophore. An *all-trans* retinal is covalently-bonded to a lysine residue through the retinal Schiff base (RSB) and stabilized by a negatively-charged counterion. The distance between the RSB and counterion is closely related to the light energy absorption. However, in heliorhodopsin-48C12 (HeR-48C12), while Glu107 acts as the counterion, E107D mutation exhibits an identical absorption spectrum to the wild-type, suggesting that the distance does not affect its absorption spectra. Here we present the 2.6 Å resolution crystal structure of the *Thermoplasmatales* archaeon HeR E108D mutant, which also has an identical absorption spectrum to the wild-type. The structure revealed that D108 does not form a hydrogen bond with the RSB, and its counterion interaction becomes weaker. Alternatively, serine cluster, S78, S112, and S238 form a distinct interaction network around the RSB. The absorption spectra of the E to D and S to A double mutants suggested that S112 influences the spectral shift by compensating for the weaker counterion interaction. Our structural and spectral studies have revealed the unique spectral shift mechanism of HeR and clarified the physicochemical properties of HeRs.

## Introduction

Heliorhodopsins (HeRs) are a family of microbial rhodopsins that was recently discovered by functional metagenomics^1^. HeRs share distant sequence identity with the type-1 (microbial) rhodopsins. The first discovered HeR was HeR-48C12, found in an actinobacterial fosmid from the freshwater Lake Kinneret and classified as a bacterial HeR. Genomic studies have led to the discovery of more than 500 rhodopsin genes in bacteria, archaea, eukaryotes, and giant viruses. HeRs have an all*-trans* retinal chromophore and undergo a photocycle involving K, M, and O intermediates upon light absorption, in parallel with retinal isomerization and proton transfer^2,3^. However, HeRs lack pump and channel activities, unlike the typical type-1 rhodopsins, and thus their functions have remained elusive. Moreover, HeRs exhibit long-lived photoactivated states with lifetimes (τ) of over a second, suggesting that HeRs are signaling photoreceptors or photoenzymes^2,4^.

Our recent crystallographic study revealed the first structure of the HeR derived from *Thermoplasmatales archaeon* SG8-52-1 (*T*aHeR)^2^. The transmembrane region of *T*aHeR consists of seven transmembrane helices, as in type-1 rhodopsins, but it is in an inverted orientation relative to them. *T*aHeR forms a stable dimer, and its interface comprises transmembrane helix (TM) 4, TM5, and two β-sheets in the ECL1. An all*-trans* retinal is covalently bound to a lysine, forming the retinal Schiff base (RSB), which is stabilized by a single counterion, E108. A linear hydrophobic pocket accommodates the retinal configuration and isomerization. Overall, *T*aHeR harbors the retinal in a similar manner to the type-1 rhodopsins, despite its many distinct features. The recently reported structure of the bacterial HeR-48C12 revealed that these characteristics are common in the HeRs^5,6^.

The RSB is protonated in type-1 rhodopsins and HeRs, in which the high pKa is stabilized and maintained by a counterion^2,7,8^. The electrostatic interaction has been extensively studied between the positively charged retinal chromophore and the negatively charged counterion, as it is the dominant component in the regulation of light energy absorption. Most type-1 rhodopsins contain two counterions at TM3 and TM7, as in the cases of D85 and D212 in the light-driven proton-pump bacteriorhodopsin (BR), and E123 and D253 in channelrhodopsin 2 from *Chlamydomonas reinhardtii* (CrChR2)^9,10^. By contrast, HeRs possess a single counterion in TM3: E108 in *T*aHeR and E107 in HeR-48C12. A previous mutation study of HeR-48C12 revealed anion binding to the E107A and E107Q mutants, but not to the wild-type and E107D mutant^8^. While the results supported E107 as the counterion, a puzzling result was obtained: identical absorption spectra were observed for the wild-type and E107D mutant^8,11^. It is well known that stronger or weaker electrostatic interactions with the counterion cause a spectral blue- or red-shift, respectively, in both type-1 and -2 rhodopsins^7^. In the case of light-driven proton pumps, the D-to-E mutants cause 28 nm, 8 nm, and 23 nm blue-shifts for BR^12^, *Gloeobacter* rhodopsin (GR)^13^, and *Acetabularia* rhodopsin I (AcetR1)^14^, respectively. By contrast, the E-to-D mutants of CrChR2 and bovine rhodopsin (type-2 rhodopsin) cause 16 nm^15^ and 10 nm^16^ red-shifts, respectively. In general, the D-to-E and E-to-D mutations cause spectral blue and red shifts, respectively. However, the absorption spectrum of the E107D mutant of HeR 48C12 is identical to the wild-type spectrum. This puzzling result suggests that either the counterion does not dominantly regulate energy absorption, or the other residues around the RSB compensate for the weaker counterion interaction.

## Material and Methods

### Determination of λ_max_ of *T*aHeR WT and mutants by hydroxylamine bleaching

#### Gene Preparation and protein Expression

The codon-optimized full-length *T. archaeon* HeR gene (GenBank ID: KYK26602.1) containing an N-terminal histidine-tag was chemically synthesized (GenScript) and subcloned into the pET21a (+)-vector, as reported previously. For mutagenesis, a QuikChange site-directed mutagenesis kit (Stratagene) was used, according to the standard protocol. After sequence confirmation, these constructs were used to transform *E. coli* strain C43 (DE3), and protein expression was induced by 1.0 mM isopropyl β-D-thiogalactopyranoside (IPTG) for 4 h at 37 °C, in the presence of 10 μM all-trans-retinal (Sigma-Aldrich).

#### Determination of Absorption Maxima without Purification

A 4 mL culture of *E. coli* expressing the desired rhodopsin was centrifuged, and the pellet was resuspended in 3 mL of buffer, containing 50 mM Na_2_HPO_4_, 100 mM NaCl, 2.0 mg lysozyme, and 10 mg DNase, pH 7.0. This mixture was gently agitated at room temperature for 1 hr. The mixture was then sonicated for complete cell lysis, combined with n-Dodecyl-β-D-maltoside (DDM) at an effective concentration (2%) for complete solubilization of the desired protein, and mixed at 4°C overnight. Afterwards, we added a freshly prepared hydroxylamine solution to the sample (final concentration, 500 mM) and illuminated it for 30 min with a 1 kW tungsten-halogen projector lamp (Master HILUX-HR, Rigaku) through a glass filter (Y-52, AGC Techno Glass) at wavelengths >500 nm. A hot mirror was placed in front of the projector lamp to block heat radiation. Absorption changes representing the bleaching of rhodopsins by hydroxylamine were measured with an ultraviolet-visible (UV-vis) spectrometer.

#### Crystallization

The E108D mutant of *T*aHeR was purified as described previously^2^. In brief, the E108D mutant was expressed in *E. coli*, solubilized with DDM, and purified by nickel affinity chromatography. The protein was concentrated to 40 mg ml^-1^ with a centrifugal filter device (Millipore 50 kDa MW cutoff), and frozen until crystallization.

The protein was reconstituted into monoolein at a weight ratio of 1:1.5 (protein:lipid). The protein-laden mesophase was dispensed into 96-well glass plates in 30 nl drops and overlaid with 800 nl precipitant solution, using a Gryphon robot (ARI), as described previously^31^. Crystals were grown at 20°C in precipitant solutions containing 25% PEG 350 MME, 100 mM Na-citrate, pH 5.0, and 100 mM ammonium sulfate or lithium sulfate. The crystals were harvested directly from the LCP with micromounts (MiTeGen) or LithoLoops (Protein Wave) and frozen in liquid nitrogen, without adding any extra cryoprotectant.

#### Data collection and structure determination

X-ray diffraction data were collected at the SPring-8 beamline BL32XU with an EIGER X 9M detector (Dectris), using a wavelength of 1.0 Å. In total, 79 small-wedge (10° per crystal) datasets using a 5×5 μm^2^ beam were collected automatically, using the ZOO syste^32^. The collected images were processed using KAMO^33^ with XDS^34^, and 52 datasets were indexed with consistent unit cell parameters. After correlation coefficient-based clustering using the intensities, followed by merging using XSCALE^34^ with outlier rejections implemented in KAMO, a small cluster consisting of 12 datasets was selected for the following analyses, because it gave a small inner-shell *R*_meas_ and high outer-shell CC_1/2_. The E108D structure was determined by molecular replacement with PHASER^35^, using the wild-type *T*aHeR structure (PDB code: 6IS6)^2^ as the template. Subsequently, the model was rebuilt and refined with COOT^36^ and REFMAC5^37^. The final model of the E108D mutant of *T*aHeR contained residues 4-253 of *T*aHeR, 14 monoolein molecules, and 33 water molecules. Figures were prepared with CueMol (http://www.cuemol.org/ja/).

## Results

### Structural determination of the E108D mutant of *T*aHeR

To investigate the effect of the E-to-D mutation in another heliorhodopsin, we measured the absorption spectra of the wild-type and E108D mutant of *T*aHeR by bleaching the retinal chromophore with hydroxylamine. The wild-type and E108D mutant exhibited identical absorption spectra, with a maximum absorption wavelength (λmax) at 542 nm (Fig. 1a, b). These data showed that the E to D counterion mutation does not cause a spectral shift, as in HeR-48C12, although the archaeal heliorhodopsin *T*aHeR and bacterial HeR-48C12 share relatively low sequence identity (43%). These observations suggest that the absence of a spectral shift by the counterion mutation E to D is a common feature in bacterial and archaeal HeRs.

**Fig. 1.**
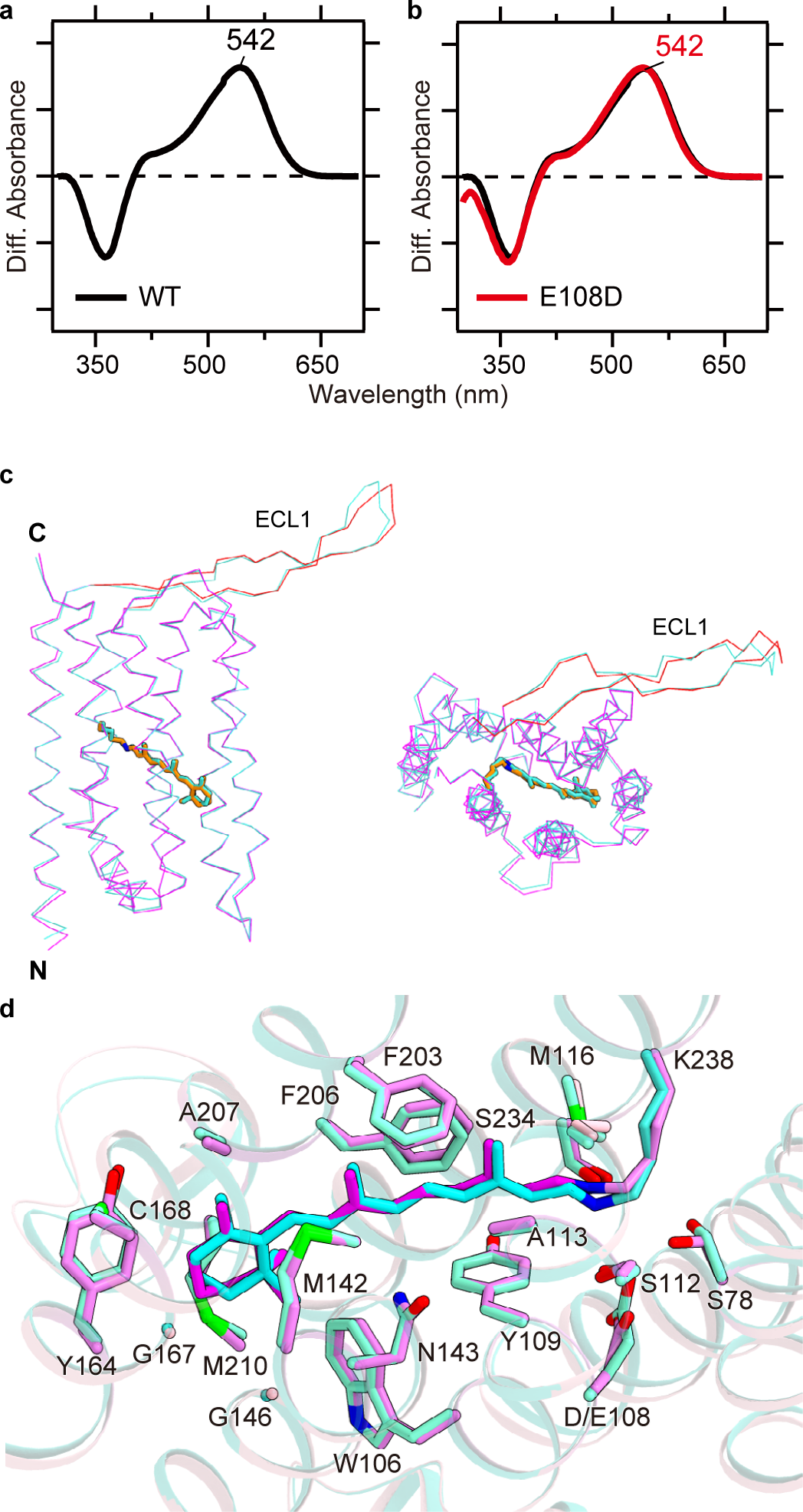
Overall structure of the *Ta*HeR E108D mutant. **a, b**, Light-induced difference absorption spectra of the WT (black curves) and the E108D mutant (red curves) of *T*aHeR in the presence of 500 mM hydroxylamine. Positive and negative signals show the spectra before and after illumination, corresponding to those of the rhodopsin and retinal oxime, respectively. **c**, Superimposed structures of the wild-type (Protein Data Bank (PDB) code: 6IS6)^2^ and E108D mutant of *T*aHeR, colored dark turquoise and magenta, respectively. **d**, Comparison of the retinal binding sites in the wild-type and E108D mutant of *T*aHeR.

To understand why the E to D mutation does not cause a spectral shift, we determined the 2.6 Å resolution crystal structure of the E108D mutant of *T*aHeR (Supplementary Fig. 1 and Supplementary Table 1). The E108D structure comprises seven transmembrane helices, a long β-sheet in ECL1, and a covalently bound *all-trans* retinal at K238 (Fig. 1c). While the crystallizing conditions and crystal packing of the E108D mutant differed from those of the wild-type (Supplementary Table 1 and Methods), it forms a similar dimer with the symmetric protomers (Supplementary Fig. 2a, b). The E108D structure superimposes well with the wild-type structure (0.4 Å root mean square deviation of Cα atoms) (Fig. 1c), indicating that the E108D mutation does not affect the overall conformation.

### Structural effect of the E108D mutation

To evaluate the effect of the E108D mutation, we compare the retinal binding pockets in the wild-type and E108D structures. The rotamers of the hydrophobic residues in the retinal binding site are almost identical (Fig. 1d). By contrast, the counterion mutation E108D alters the polar interaction network around the RSB. In the wild-type structure, the distance between the RSB and counterion carboxylate is 3.5 Å (Fig. 2a). Thus, they form a hydrogen bond in addition to the electrostatic interactions, as also supported by a previous resonance Raman analysis^3^. In the E108D structure, this distance becomes longer (4.9 Å), and thus the counterion can no longer form a hydrogen bond with the RSB (Fig. 2b and Supplementary Fig. 3). We could not observe a density corresponding to a water between the counterion and RSB in the 2.6 Å resolution structure (Supplementary Fig. 3), suggesting that they do not even form a water-mediated hydrogen-bonding interaction. As the interaction between the RSB and counterion becomes weaker, we expected the E108D mutant exhibits a blue-shifted absorption.

**Fig. 2.**
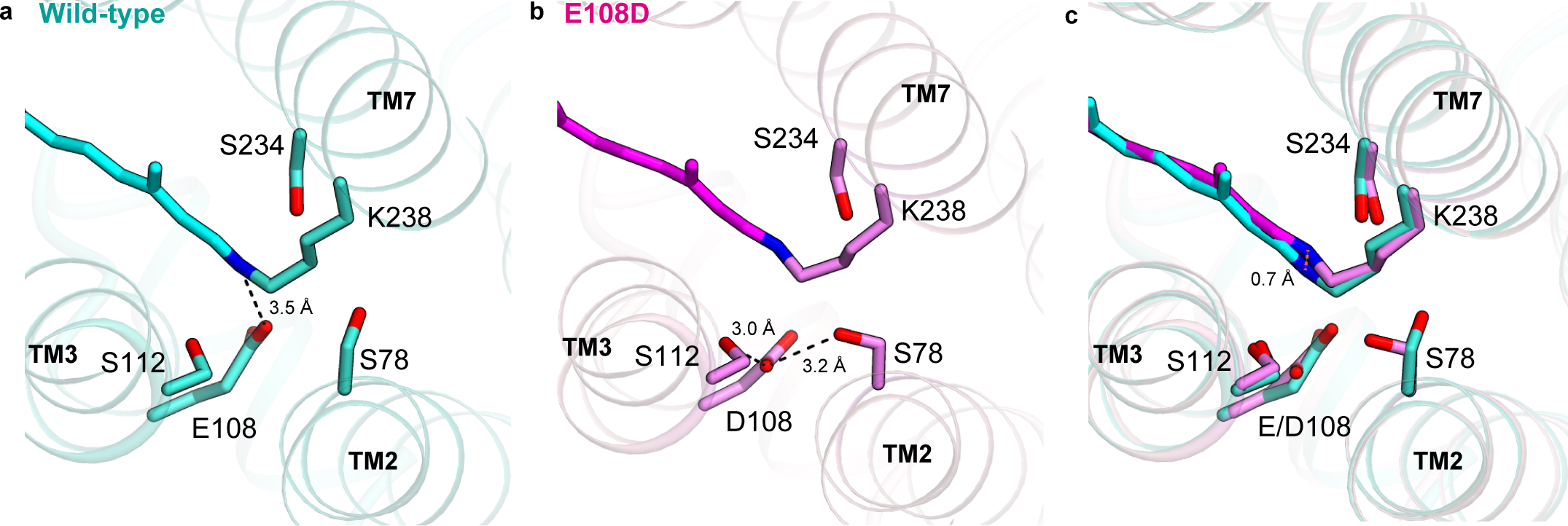
Interactions around the retinal Schiff base. **a, b**, Comparison of the interactions around the RSB and counterion in the wild-type (**a**) and E108D mutant (**b**), viewed from the extracellular side. Hydrogen-bonding interactions are indicated by black dashed lines. The water is shown as a red sphere. **c**, Overlay of **a** and **b**.

Notably, the E108D mutation rearranges the polar interaction network around the RSB-counterion complex. In the wild-type structure, the hydroxyl groups of S112 in TM3 and S234 in TM7 sandwich the RSB and form polar interactions (Fig. 2a). The counterion E108 only forms a hydrogen bond with the RSB. In the E108D structure, the distance between the hydroxyl group of S112 and the counterion is closer (3.8 Å to 3.0 Å) to allow a hydrogen-bonding interaction (Fig. 2b). Moreover, the S78 side chain in TM2 flips and also forms a hydrogen bond with the counterion (Fig. 2c). The RSB moves in the opposite direction from the counterion by 0.7 Å, and the S234 side chain also shifts slightly. Although the interaction between the counterion and RSB certainly becomes weaker, the interactions involving S78, S112, and S234 are rearranged (Supplementary Table 3) to establish additional hydrogen-bonding interactions in the E108D structure (S78-D108 and S112-D108). These serine residues are conserved in *T*aHeR and HeR-48C12. The aforementioned hydrogen-bonding interactions would compensate for the weaker counterion interaction by the E108D mutation and retain the absorption spectrum.

### Mutational analysis around the RSB

To examine the effects of these serine residues, we measured the absorption spectra of the single mutants (S78A, S112A, and S234A) and the double mutants (S78A/E108D, S112A/E108D, and S234A/E108D). If these serine residues compensate for the weaker counterion interaction, then the S to A/E108D double mutant should show red-shifted absorption as compared with that of the single S to A mutant. First, we describe the results of the S78 and S234 mutations (Fig. 3a-d). The single mutants S78A and S234A showed a 2 nm red-shifted absorption at 544 nm, as compared with that of the wild-type (Fig. 3a, c). Unexpectedly, the S78A/E108D and S234A/E108D double mutants showed maximum absorption wavelengths at 532 nm and 540 nm, which are blue-shifted by 14 and 4 nm, respectively (Fig. 3b, d). These results suggest that S78 and S234 are not associated with the lack of a spectral shift by the E108D mutation. We cannot precisely explain these blue-shifted absorptions, and especially that of the S78A/E108D double mutant. The S78A/E108D double mutation shortens their side chains and would create more space around the counterion. This extra space may allow water molecules to enter and stabilize the RSB by additional water-mediated hydrogen-bonding interactions, resulting in the blue-shifted absorption spectra.

**Fig. 3.**
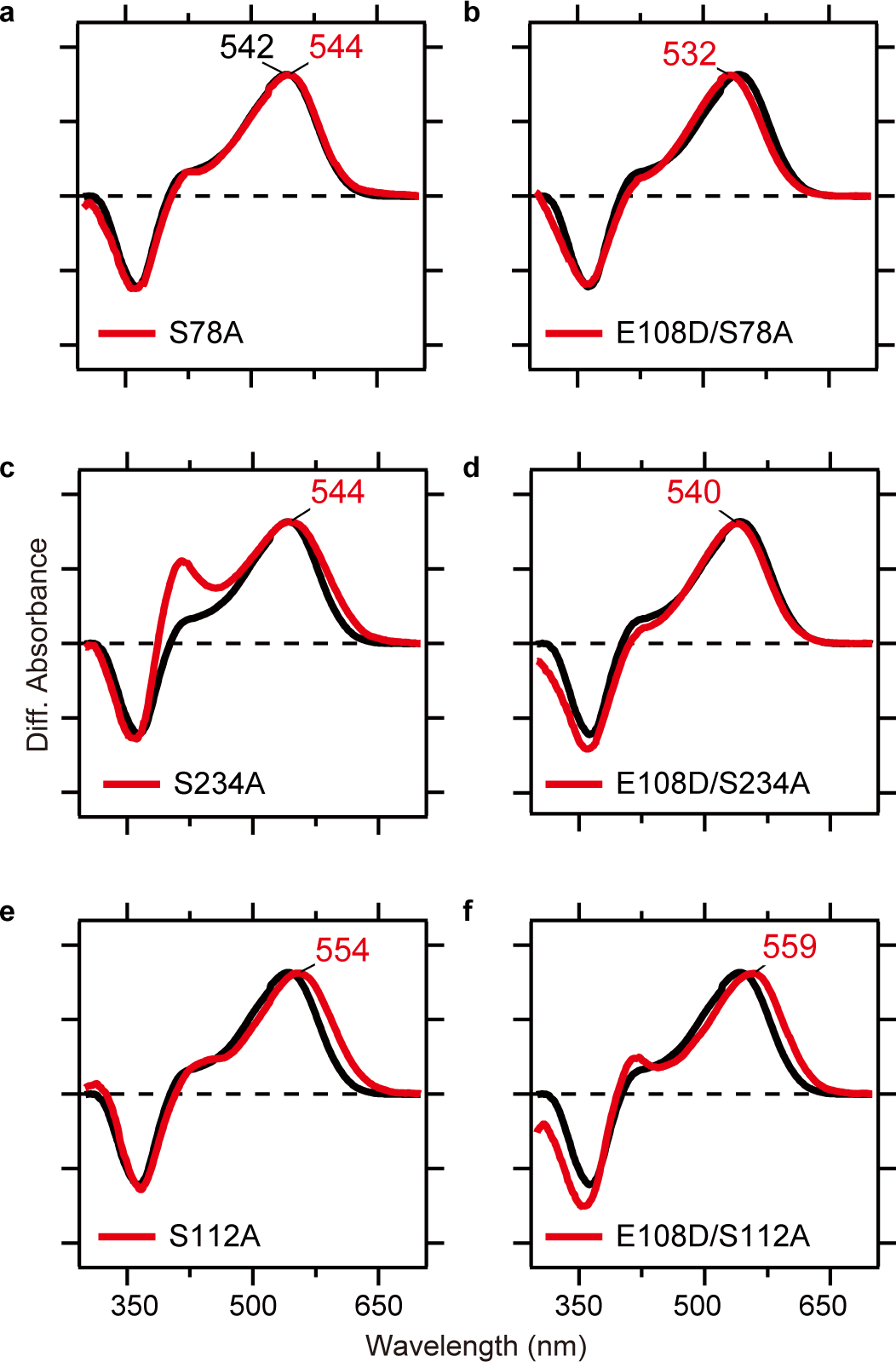
Spectroscopic analysis of *T*aHeR mutants. **a-f**, Light-induced difference absorption spectra of the WT (black curves) and mutants of the Schiff base region (red curves) of *T*aHeR in the presence of 500 mM hydroxylamine. Positive and negative signals represent the spectra before and after illumination, corresponding to those of the rhodopsin and retinal oxime, respectively.

Next, we describe the results obtained with the S112 mutant. The S112A mutant showed a 12 nm red-shifted absorption at 554 nm, as compared with that of the wild-type (Fig. 3e), indicating that it plays a critical role in stabilizing the RSB. Moreover, the maximum absorption wavelength (λmax) of the S112A/E108D double mutant was 559 nm (Fig. 3f), which is 5 nm red-shifted as compared with that of the S112A mutant. This result indicates that the E108D mutation causes the spectral red shift in the *T*aHeR S112A mutant. Therefore, S112 compensates for the weaker counterion interaction by the E108D mutation and maintains the absorption spectra. The polar interactions of S112 with the RSB-counterion complex would be essential for this effect (Fig. 2b).

S112 is located just one helical turn above the counterion (E108) and conserved among most HeRs (97%). The distances between the hydroxyl group of the serine and RSB are 3.0 Å and 3.2 Å in *T*aHeR and HeR-48C12, respectively^2,5^ (Fig. 4a, b). By contrast, the equivalent residue is conserved as a threonine in most type-1 rhodopsins. The distance between the hydroxyl group of the threonine and RSB is about 3.7 Å in the representative type-1 rhodopsins (BR, CrChR2, and rhodopsin phosphodiesterase)^9,10,17,18^ (Fig. 4c-e), which is longer than those in the HeRs. Marti *et al*. reported largely blue-shifted absorption spectra for the BR T89A mutant, when expressed in *E. coli*^19^. However, when expressed in *H. salinarum*, the native cells of BR, the T89A mutant showed similar absorption spectra to those of the wild type^20^. Moreover, Ehrenberg *et al*. reported CrChR2 T127A mutation shifted the visible absorption spectrum only slightly to the blue as compared to the wild-type^21^. These studies indicate that the threonine is not involved in stabilizing the RSB, in contrast to S112 in *T*aHeR. This difference is one of the critical factors determining whether the counterion mutation E to D causes a spectral shift.

**Fig. 4.**
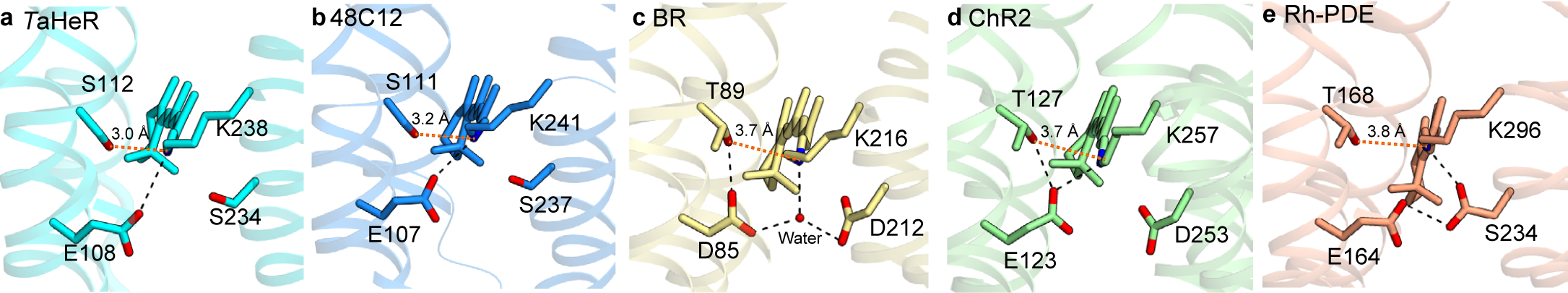
Comparison of the residues around the RSB. **a-e**, Interactions around the RSB in *T*aHeR (PDB code: 6IS6)^2^, HeR-48C12 (PDB code: 6SU3)^5^, BR (PDB code: 1C3W)^9^, CrChR2 (PDB code: 6EID)^10^, and Rh-PDE (PDB code: XXX)^17^. Hydrogen-bonding interactions are indicated by black dashed lines. The water is shown as a red sphere.

## Discussion

Our study revealed that the E to D mutation of the counterion does not alter the absorption spectrum of *T*aHeR, suggesting that this feature is common among HeRs. The crystal structure of the E108D mutant revealed that the mutation only alters the environment around the RSB. The E108D mutation certainly increases the distance between the counterion and RSB, and affects the polar interaction network between the RSB, the nearby serine residues, and the counterion. Notably, S112 and the counterion form an additional hydrogen bond in the E108D structure. The mutant analysis showed that the E108D mutation causes a 5 nm red-shift in the S112A mutant, suggesting that S112 stabilizes the RSB by the hydrogen-bonding interaction with the counterion and thus retains the absorption spectra. Our study suggests that S112 compensates for the weaker counterion interaction by the E108D mutation. This unique feature of *T*aHeR might be associated with its physiological function.

The mechanism of color tuning in type-1 and -2 rhodopsins has fascinated researchers for a long time, as it relates to our color discrimination^7,22–27^. While light energy absorption is determined by complex factors to control the energy gap between the ground and excited states of the retinal chromophore, the electrostatic interaction of retinal with the counterion(s) is the most prominent factor in color tuning. Despite the significant improvements of theoretical calculations^28–30^, full reproductions of the absorption spectra from the structures of rhodopsins by computation remain difficult. The present study has provided a theoretical challenge, where the weakened interaction of the counterion in E108D is compensated by the reorganized environment around counterion, leading to identical absorption maxima between the mutant and wild-type *T*aHeRs.

## Acknowledgements

The diffraction experiments were performed at SPring-8 BL32XU (proposal (2019B2577). We thank the members of the Nureki lab and the beamline staff at BL32XU of SPring-8 (Sayo, Japan) for technical assistance during data collection. This research was partially supported by the Platform Project for Supporting Drug Discovery and Life Science Research (Basis for Supporting Innovative Drug Discovery and Life Science Research (BINDS)) from AMED under grant number JP19am0101070 (support number 1627). This work was also supported by JSPS KAKENHI grants 16H06294 (O.N.), 17J30010, 30809421, 20K15728, 20H05437 (W.S.), 25104009, 18H03986, 19H04959 (H.K.), and by JST CREST (JPMJCR1753 to H.K.).

## Contributions

T.T. purified, crystallized, and solved the structure of the E108D mutant of *T*aHeR. M.S. performed spectroscopic analyses. K.Y. assisted with the structural determination. W.S, H.K. and O.N. supervised the research.

## Data Availability

Coordinates and structure factors have been deposited in the Protein Data Bank, under the accession number XXXX. The X-ray diffraction images are also available at the Zenodo data repository (https://doi.org/10.5281/zenodo.3871080).

## Supplementary Figures

**Supplementary Fig. 1.**
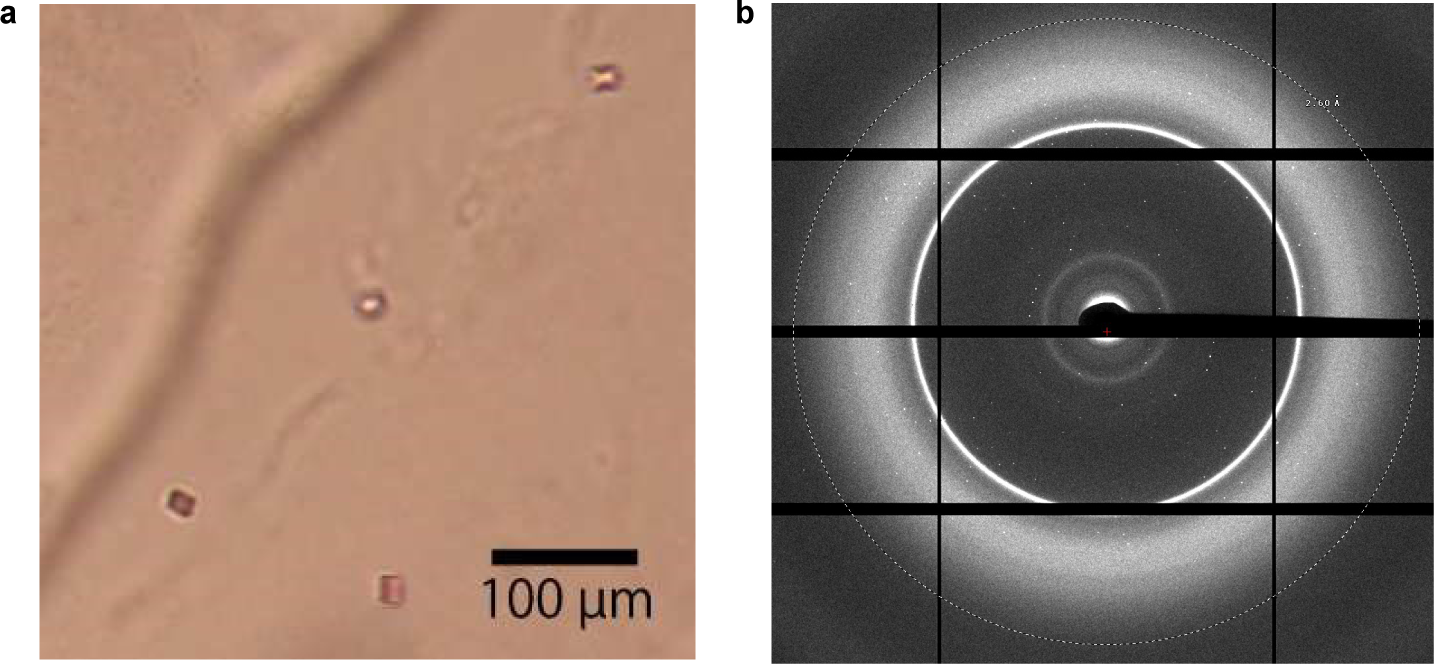
Crystallization. **a, b**, Crystals of the *T*aHeR E108D mutant (**a**) and its diffraction image (**b**).

**Supplementary Fig. 2.**
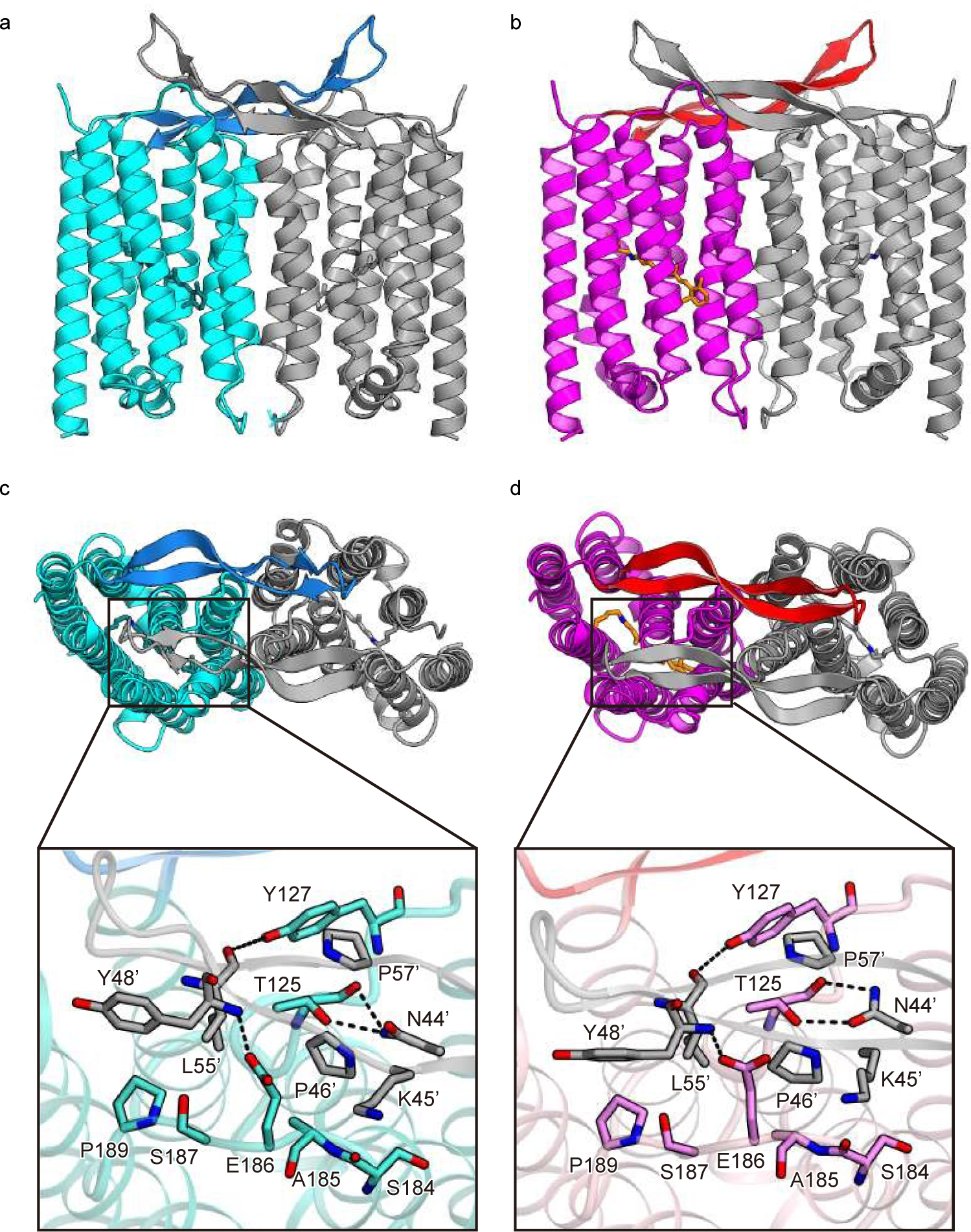
Dimer interface. **a, b**, Dimer interfaces in the *T*aHeR WT (**a**) and E108D (**b**) structures. **c, d**, Panels showing the interactions between the transmembrane region and the ECL1 in the symmetric protomers of *T*aHeR WT (**c**) and the E108D mutant (**d**). Hydrogen bonding interactions are indicated by black dashed lines.

**Supplementary Fig. 3.**
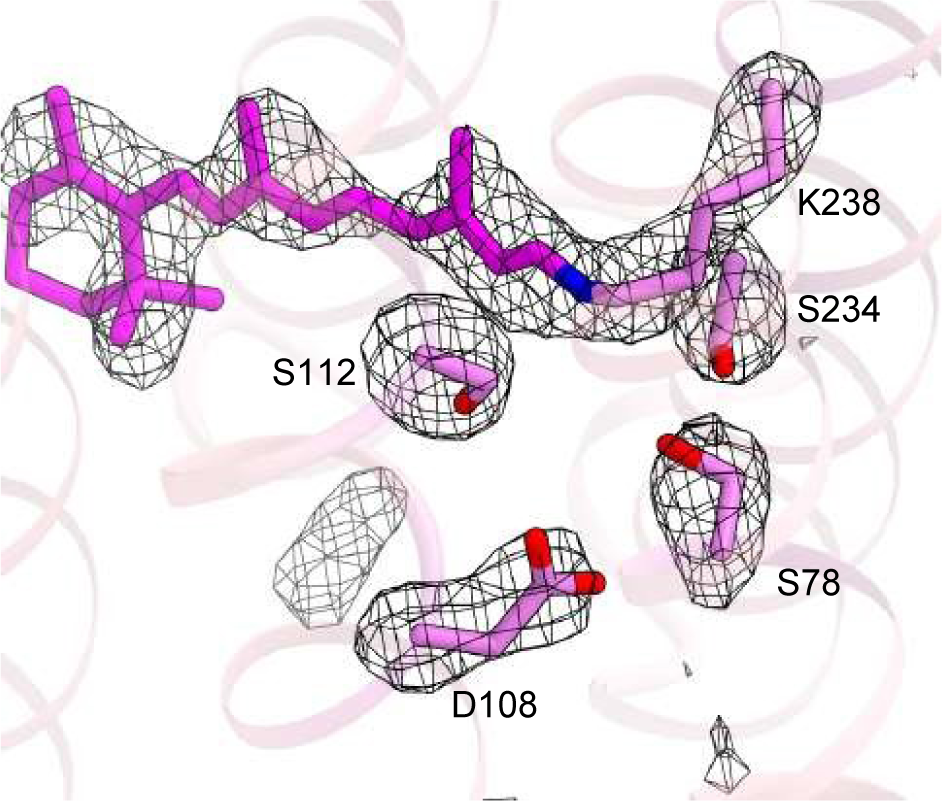
Electron density around the counterion. The polder omit map (*mF*_*o*_-*DF*_*c*_)^38^ contoured at 3.2σ, calculated by omitting the retinal and side chains of S78, D108, S112, K234, and S238.

**Supplementary Table 1.** Data collection and refinement statistics. *Values in parentheses are for the highest-resolution shell.

|  | Native |
| --- | --- |
| **Data collection** |  |
| Space group | *P4<sub>2</sub>2<sub>1</sub>2* |
| Cell dimensions |  |
| a=b, c (Å) | 72.9, 115.0 |
| Resolution (Å)<sup>a</sup> | 30-2.60 (2.75-2.60) |
| R<sub>meas</sub><sup>a</sup> | 0.380 (1.806) |
| <I/σ(I)><sup>a</sup> | 5.36 (1.04) |
| CC<sub>1/2</sub><sup>a</sup> | 0.982 (0.484) |
| Completeness (%)<sup>a</sup> | 99.1 (98.6) |
| Redundancy<sup>a</sup> | 7.71 (7.60) |
| **Refinement** |  |
| Resolution (Å) | 30-2.6 |
| No. unique reflections | 9975 |
| R<sub>work</sub>/R<sub>free</sub> | 0.2215/0.2834 |
| No. atoms |  |
| Protein | 2007 |
| Ligand | 250 |
| Water | 15 |
| Averaged B-factors (Å<sup>2</sup>) |  |
| Protein | 38.9 |
| Ligand | 62.2 |
| Water | 33.8 |
| R.m.s. deviations from ideal |  |
| Bond lengths (Å) | 0.0049 |
| Bond angles (°) | 1.3595 |
| Ramachandran plot |  |
| Favored (%) | 97.18 |
| Allowed (%) | 2.82 |
| Outlier (%) | 0 |

<sup>a</sup>Values in parentheses are for the highest-resolution shell.

**Supplementary Table 2.**

| Distance from RSB (Å) | WT | E108D |
| --- | --- | --- |
| E/D108 | 3.5 | 4.9 |
| S78 | 4.6 | 3.8 |
| S112 | 3.0 | 3.7 |
| S234 | 3.2 | 3.6 |

